# How signals of calcium ions initiate the beats of cilia and flagella

**DOI:** 10.1101/585034

**Authors:** M. V. Satarić, T. Nemeš, D. Sekulić, J. A. Tuszynski

## Abstract

Cilia and flagella are cell organelles serving basic roles in cellular motility. Ciliary movement is performed by a sweeping-like repeated bending motion, which gives rise to a self-propagating “ciliary beat”. The hallmark structure in cilia is the axoneme, a stable architecture of microtubule doublets. The motion of axoneme is powered by the axonemal dynein motor family powered by ATP hydrolysis. It is still unclear how the organized beat of cilium and flagella emerges from the combined action of hundreds of dynein molecules. It has been hypothesized that such coordination is mediated by mechanical stress due to transverse, radial or sliding deformations. The beating asymmetry is crucial for airway ciliary function and it requires tubulin glutamination a unique posttranslational modification of C-termini of constituent microtubules that is highly abundant in cilia and flagella. The exact role of tubulin glutamination in ciliary or flagellar function is still unclear. Here we examine the role of calcium (Ca^2+^) ions based on the experimental evidences that the flagellar asymmetry can be increased due to the entry of extracellular Ca^2+^ through, for example, nimodipine-sensitive pathway located in the flagella. We propose a new scenario based on the polyelectrolyte properties of cellular microtubules (MTs) such that dynamic influx of Ca^2+^ ions provides the initiation and synchronization of dynein sliding along microtubules. We also point out the possible interplay between tubulin polyglutaminated C-termini and localized pulses of Ca^2+^ ions along microtubules.

## INTRODUCTION

Several micro-organisms and cells swim in a viscous medium due to the active rhythmic wave-like motion of cilia and flagella. Motile cilia and flagella are capable of complex finely coordinated movements and have subtle versatile roles in embryonic development, fertilization and function of epithelia (1). Cilia are important not only for cell motility but also for their sweeping-like action that is, for instance, seen in the trachea where they perform the clearing mucus out of lungs. On the other hand, the flagellum forms the “tail” on sperm cell that propels it in order to swim. Both cilia and flagella contain an axoneme, which consist of a cylindrical arrangement of nine doublets of parallel microtubules and a pair of microtubules in the cylinder’s center called the central apparatus(see Figure 1(a)). Additionally, Ikegami et al. (2), reported that the beating asymmetry is crucial for airway ciliary function and requires tubulin glutamination, which is a unique post-translational modification of C-termini of constituent microtubules that is highly abundant in cilia and flagella. There is a significant number of associated proteins, such as nexin which provides elastic tangential links between the microtubule doublets, and radially distributed links named radial spokes (3).

**Figure 1:**
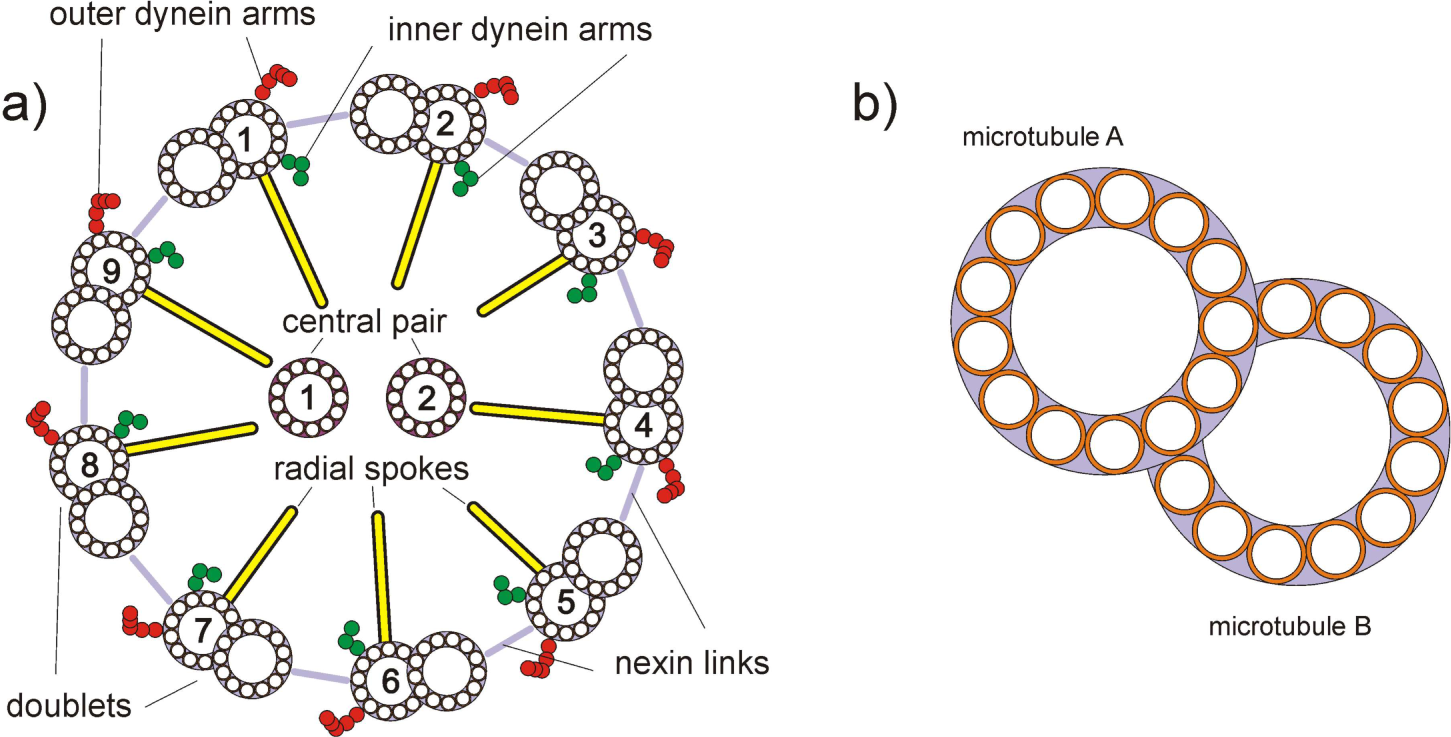
a) Schematic of an axonemal cross section of radius *a*_0_. The nine doublets are numbered. b) Geometry of axonemal doublet.

Each microtubule doublet within an axoneme is connected to its nearest-neighbors by cross-linkers such as nexin. These nexins provide a resistance to the relative sliding of doublets and the change in the spacing between them. The inter-spacing of doublets is approximately 30 nm, which is similar in size to the diameter of a microtubule, namely 25 nm. The doublets are composed of one A-microtubule and one B-microtubule. The A-microtubule has 13 parallel protofilaments while the B-microtubule has only 10 protofilaments, Figure 1(b).

Radial spokes keep the diameter of the axoneme at approximately 200 nm in length maintaining the cross-section of the axoneme. Microtubule doublets are dynamically connected by a large number of dynein molecular motors, which produce active forces in terms of chemical energy provided by ATP hydrolysis. Dynein motors generate force and torques that slide and bend the constituent microtubule doublets. In order to create regular wave-like movements of the axoneme, the action of dynein motors should be properly coordinated. All these elements (nexin, radial spokes and dynein motors) are loacted periodically along the long axis of axoneme cylinder with a period of approximately 96 nm.

Dyneins are rigidly attached to the A-microtubules and their stalks are briefly in contact with the adjacent B-microtubule during the power stroke process. For example, chlamydomonas axoneme contains 14 different types of dyneins and has a total of 10^4^ of these motors over its length of about 10 *µm* (4).

Bending of the axoneme originates from the imbalance of dynein motor forces on the opposite sides of the bending plane (5). The axoneme is equipped with two qualitatively different, largely independent systems for bending; one just consisting of outer-arm dyneins and the other involving inner-arm dyneins cooperating with the central-pair and radial spokes.

There is a strikingly large difference in structure and organization between outer- and inner-arm dyneins within the axoneme. Chlamydomonas axoneme has 12 outer dyneins (grouped in 4 outer dynein-arms) and 8 inner dyneins (2 dimetric I_1_ dynein, and 6 single headed inner dynein-arms) per each 96 nm repeat.

During their power stroke dyneins produce forces that tend to slide the axonemal doublets with respect to each other, thus regulating the beat pattern of the axoneme. It is believed that the ciliary and flagellar beat is enabled by alternating episodes of activation of opposing sets of dynein as a self-organized process in such a way that dyneins regulate the beat, and conversely, the beat tunes the dyneins.

When activated dyneins on one side of the axoneme win the tug-of-war, this leads to relative motion between the microtubule doublets (6). Passive nexin linkers constrain sliding and convert it into bending. So far the following three mechanism of dynein regulation within axoneme have been proposed:

a. Sliding control, which produces a load force that triggers dynein detachment (7). This mechanism has shown good fit to experimental data for beating of the bull sperm (8).
b. Normal force control, or geometric clutch control, where the increase in the distance between doublets creates a normal force that tends to detach dyneins (9).
c. Curvature control, where the switching of dyneins is regulated by the curvature of the axoneme, (10). However, the problem is due to the small size of dynein compared to the curvatures in the axoneme, which makes such geometrical sensing unlikely (11). We speculate that C-termini of MTs can be sensitive to curvature.

Major question about cilia and flagella beating initiation events still remain unsolved, namely (5):

1. What determines when and how a dynein sliding initiation events occur?
2. What mechanism allows sliding to be initiated so synchronously over an extended region?

Interestingly, the concepts dealing with the above-mentioned mechanism of dynein regulation do not include the influence of calcium signaling. On the other hand, there is abundant experimental evidence about importance and even essential roles that Ca^2+^ ions play in the dynamics of cilia and flagella (12–14).

This article attempts to address how calcium localized signals can initiate dynein sliding within an axoneme. This promising scenario relies on the polyelectrolyte properties of constituent microtubules. We previously elaborated (15–20) on the model of localized ionic pulses propagating along cellular microtubules based on the concept of a nonlinear electric transmission line. The section below describes the following aspects of the subject

1. Short revision of the concept of polyelectrolyte-induced localized Ca^2+^ pulses propagating along microtubules.
2. Short introduction to the basic differential equations of flagella dynamics within the curvature-regulated beating model.
3. Implementation of Ca^2+^ pulses in initiation of dynein sliding in the axoneme.
4. Qualitative description of possible roles of post-translational polyglutamylation of tubulin’s C-terminals in axoneme dynamics is also discussed.

### MICROTUBULES (MTS) AS POLYELECTROLYTES ACTING AS ELECTRIC TRANSMISSION LINES FOR Ca^2+^ IONS

Relying on a few experimental assays (21, 22), which indicated that the presence of MTs in cytosolic solution increases its ionic conduction, we earlier established an original model which predicts the propagation of signaling ionic impulses along MTs in the form of solitons (15, 16, 19).

We here briefly reexamine this concept in order to apply it for MTs contained in the axoneme of cilia and flagella. MTs are mostly negatively charged on their outer surface due to the numerous amino acids forming tubulin protein that have negatively charged residues under physiological conditions.

Each tubulin monomer of the MT lattice has a short C-terminal helix H12 followed by highly acidic amino acid sequence protruding out of the MT surface called a C-terminal tail (CTT), with a length 𝓁_*CTT*_ = 4.5*nm* when completely outstretched. These CTTs are very essential for the interactions of MTs with motor proteins and other MT-associated proteins (MAPs). More details about CTTs can be found in (23).

At neutral pH the very negative charge on a CTT causes it to remain extended due to electrostatic repulsion. Under more acidic conditions the negative charges of the amino acids within CTT become partially neutralized by counter-ions loosely condensed on them. This effect allows CTTs to partially shrink and fold. This is the largest conformational effect in MTs and it plays an important role in the framework of the model presented here.

The *α-CTT* is 19 amino acid long and has 10 negatively charged residues in the absence of post-translational modifications. The *β-CTT* has 9 negatively charged residues (E,A). It should be stressed here that post-translational modifications in MTs are mostly concentrated on either *α* or *β*-CTTs (2, 24–28).

This means that in the case of polyglutamination the negative charge of CTTs is markedly increased thus influencing the electric capacity of the associated MTs in the context of the model developed here.

Keeping in mind that MTs together with their CTTs are mostly negatively charged with high enough charge density, so that they can be refereed as polyelectrolyte polymers in the context of Manning’s theory (29, 30). For example, for CTTs without post-translational modifications the linear spacing of elementary charges along a CTT is on the order of

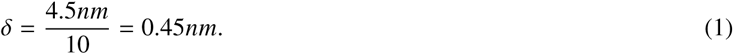

This value Bjerrum length 𝓁_*B*_ determined by the condition:

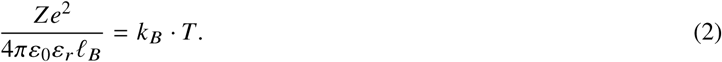

Taking *Z* = 2 for Ca^2+^, *ε*_0_ = 8.85 *·* 10^*-*12^ *F/m, ε*_*r*_ = 80, *k*_*B*_ = 1.38 *·* 10^*-*23^ *J/K, e* = 1.6 *·* 10^*-*19^ and *T* = 310*K*, one obtains

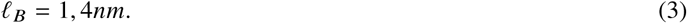

This yields a characteristic parameter *ζ* as

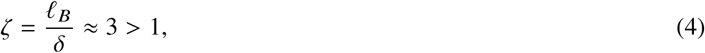

which satisfies the necessary condition for MT polymer to be considered a polyelectrolyte. The basic idea here relies on the fact that negatively charged surface of a MT and the associated CTTs attracts positively charged counterions from cytosol. This is mainly in terms of divalent cations such as Ca^2+^ and Mg^2+^, bringing them very close to the surface and forming an approximately cylindrical condensed “ionic cloud” as shown in Figure 2, with a thickness on the order of 1 nm and an inner radius of MT and CTT, respectively.

**Figure 2:**
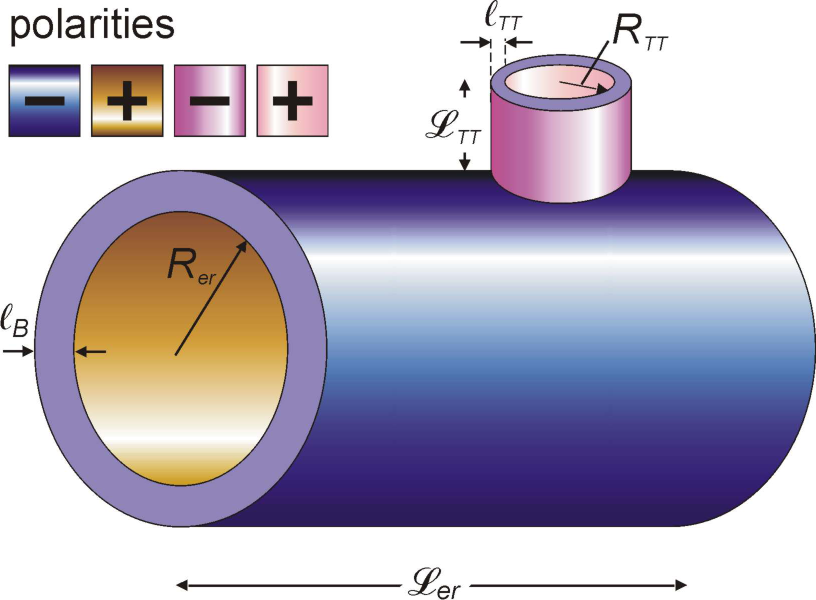
Elementary ring of an MT with characteristic cylindrical capacitors and corresponding dimensions.

These Ca^2+^ ions are not bound but allowed to slide along MT. Around this “ionic clod” there is a layer depleted of ions of both signs with a thickness equal to the Bjerrum length 𝓁_*B*_. Consequently, we can establish the elementary ring of an MT consisting of 13 parallel tubulin dimers with the associated 26 CTTs. This plays the role of a tiny cylindrical capacitor capable of storing and transporting an “ionic cloud” of attracted cations including Ca^2+^ons along the length of an MT. By calculating the electrical capacity of an elementary ring of an MT with the length ℒ_*er*_ = 8*nm* and radius *R*_*er*_ = 12.5*nm*, using the expression

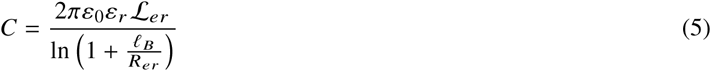

and including 13*×*2 extended CTTs with parameters ℒ_*TT*_ = 4.5*nm* and *R*_*TT*_ = 0.5*nm*, we obtain the total capacitance of this elementary capacitor to be the sum of a contribution due to the MT ring and 26 CTTs (16).

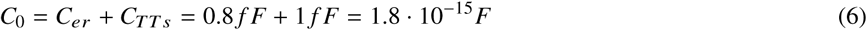

It is clear that CTTs play a dominant role in capacitance, even in the case without polyglutamination. This means that in the case of polyglutaminated MTs in cilia and flagella, the additional negative charges will increase the associated capacitance of *C*_*TTs*_. This circumstance has influence on the amplitude and speed of the “ionic clouds” surrounding MTs within axoneme, Figure 3.

**Figure 3:**
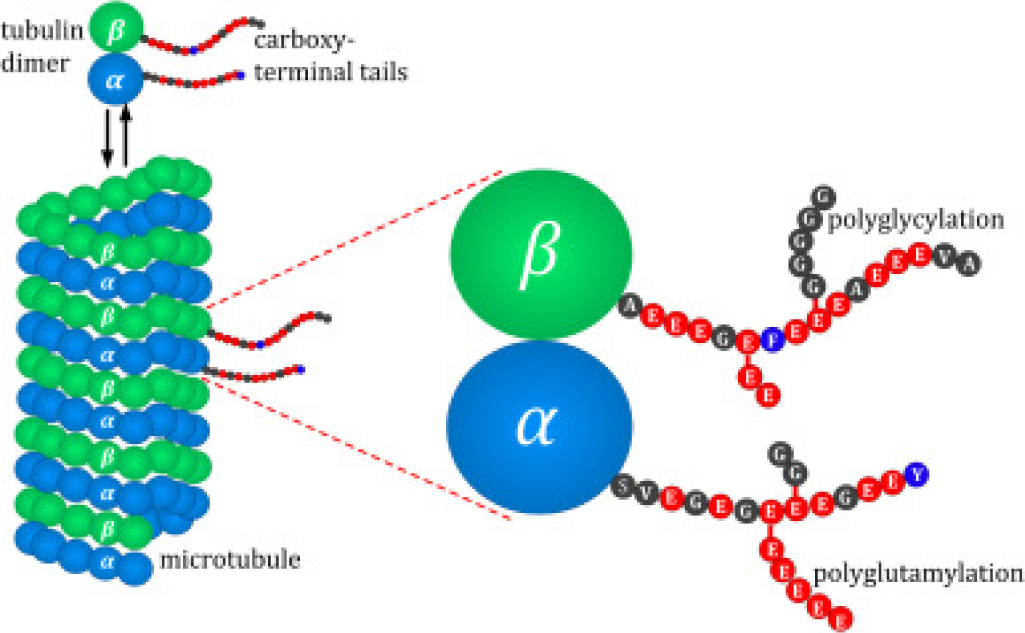
Schematic representation of the microtubule structure and distribution of different post–translational modifications of tubulin heterodimers with respect to their position in the microtubule lattice.

Above, we have elaborated on three slightly different possible mechanismss involving MTs as nonlinear transmission lines for ionic propagation (15, 16, 19). The non-linearity originates from nonlinear capacitance of elementary rings, which chiefly arises due to the flexible topology of CTTs. The nonlinear charge accumulation for the *n*-th elementary ring reads:

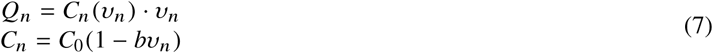

with *b*[*υ*^*-*1^] being the parameter of non linearity. Otherwise, we also considered nonlinear negative differential resistance of nano pores within each elementary ring with the associated variable admittance *G*_*n*_ containing time derivative of voltage:

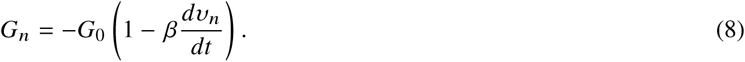

It has also been explained earlier why the inductance of MTs seen as the transmission line can be safely ignored (15). Based on the experimental findings of Minoura and Muto (22) we estimated the resistivity of an elementary ring to be on the order of 1*G*Ω:

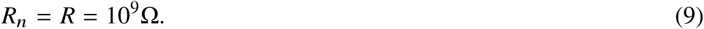

This has been established based on the effective circuit diagram in the context of Kirchhoff’s laws constructed for a repetitive arrangement of elementary rings of an MT considered as a ladder-like line, Figure 4.

**Figure 4:**
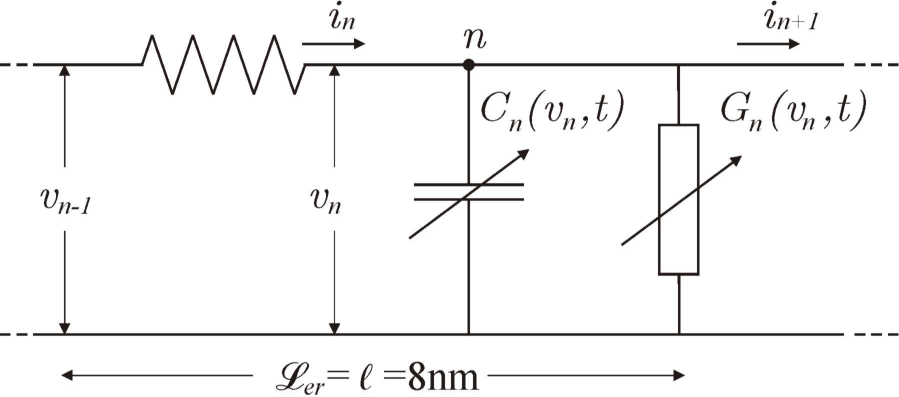
An effective circuit diagram for the *n*-th elementary unit of a transmission line with characteristic elements for Kirchhoff’s laws. 𝓁= 8*nm* is the length of a tubulin dimer equal to the length of an elementary ring of the circuit.

**Figure 5:**
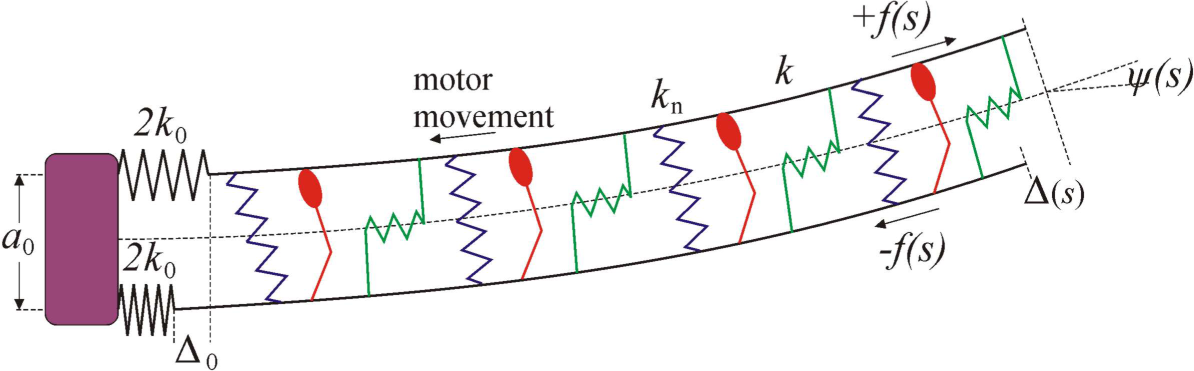
Schematic representation of two adjacent MT doublets bent by the action of a collection of dynein motors.

**Figure 6:**
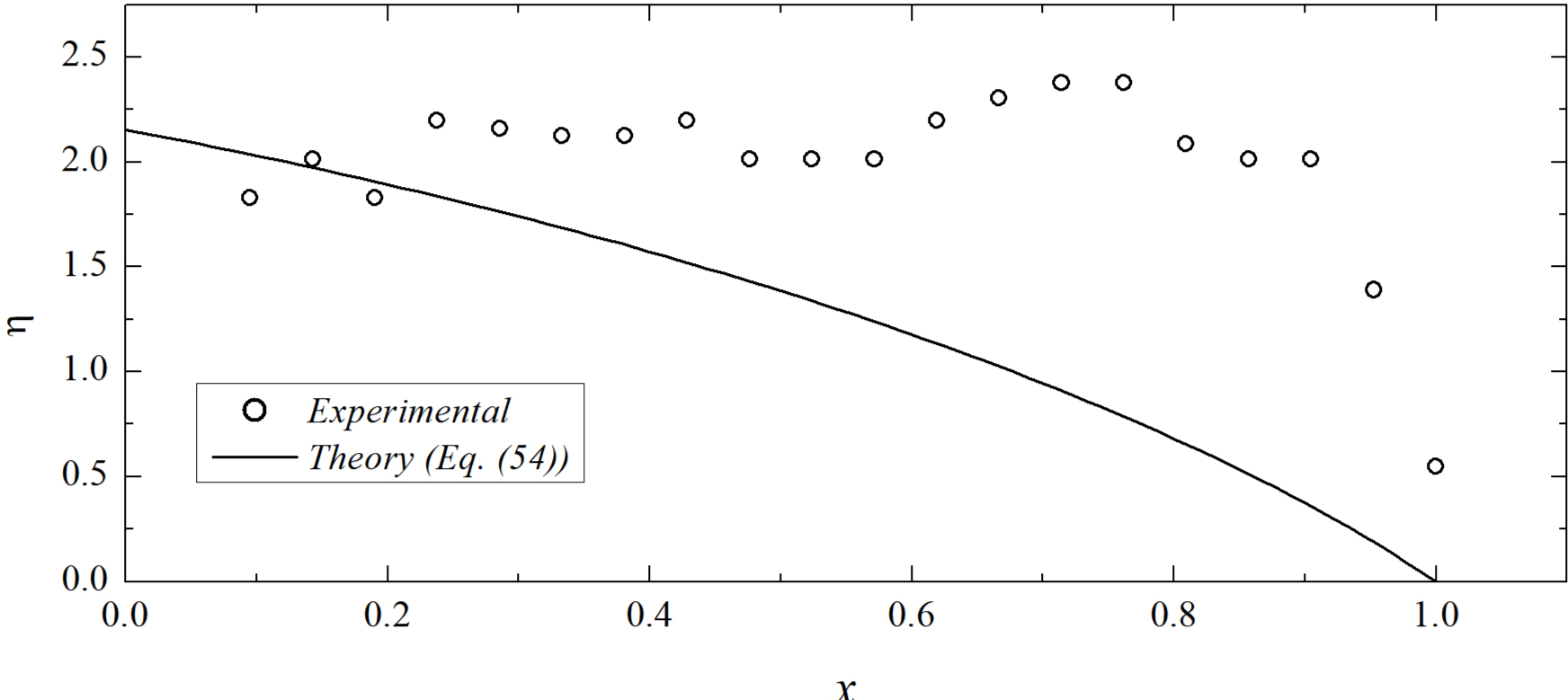
Comparison of the normalized observed filament shape (31) to the theoretical curve Eq. 54.

The Kirchhoff’s laws applied to this case here read:

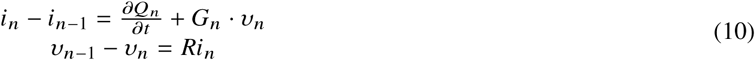

Then, it follows that the introduction of an auxiliary variable *u*_*n*_ through local voltage *υ*_*n*_ and a corresponding current *i*_*n*_, involving the impedance *Z* as

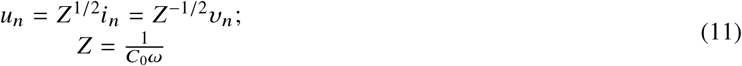

Expanding the discrete difference equations, Eq. 10 into the continuum limit of a Taylor series in terms of a small spatial parameter 𝓁= 8*nm*, and we obtain:

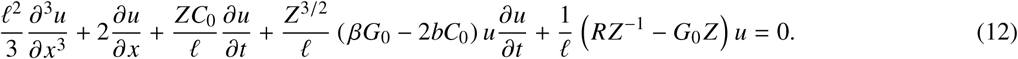

We then simplified the equation by introducing dimensionless variables in Eq. 12 using the characteristic discharging time *T*_0_ and speed *v*_0_:

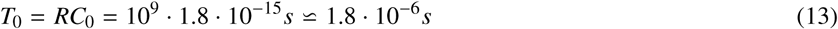

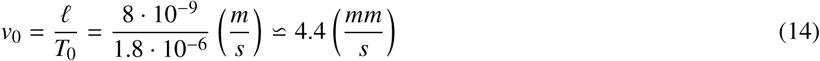

and adopting the form of progressive traveling wave with actual speed *v* and relative speed 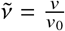

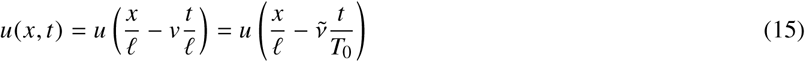

and using dimensionless unified space-time variable *ξ*

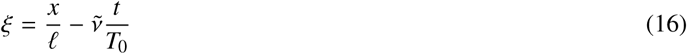

we transform the partial differential equation Eq. 12, into the corresponding ordinary differential equation as follows:

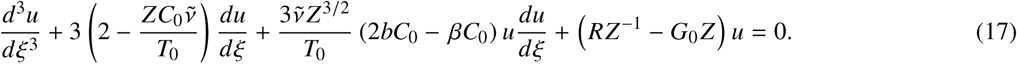

Introducing the dimensionless current 𝔍

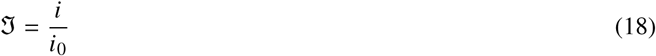

where *i*_0_ denotes the peak value, yields

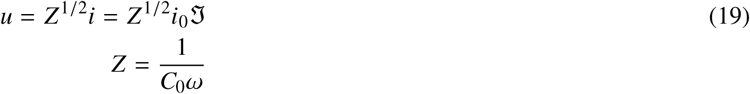

so that the Eq. 17 eventually takes the compact form:

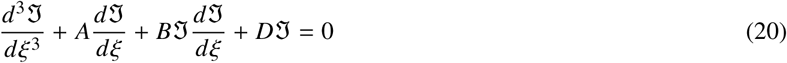

with the suitable set of abbreviations:

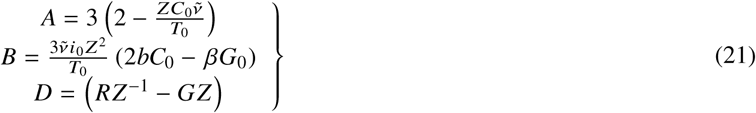

In order to solve Eq. 20 we use a new auxiliary variable *y*(*ξ*) by the definition

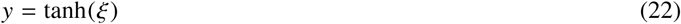

After a straightforward procedure given in Appendix the localized bell-shaped soliton can be obtained with the following functional form

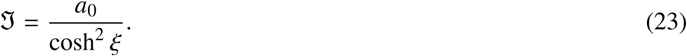

The restrictions arising from the substitution of Eq. 23 into Eq. 20 lead to the following three relations between model parameters:

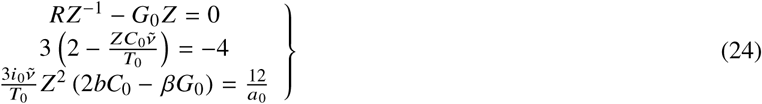

The first equation allows us to estimate the value of nanopore conductance. Taking *Z* = 2.8 *·* 10^10^Ω one obtains

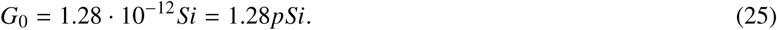

The second equation enables an assessment of the relative speed 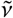 of the ionic pulse

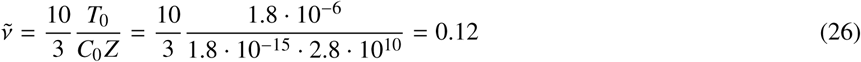

Since 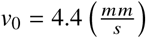 we conclude that the ionic pulse propagates with a speed of

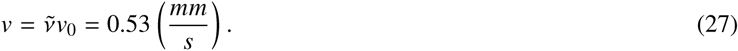

Hence, the length of a chlamydomonas *L* = 12*µm* can be traversed by this pulse during a time interval of

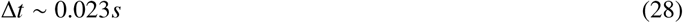

while the beating frequency of chlamydomonas is on the order of 50*Hz*, (11). The time interval in Eq. 28 would correspond to a fact that a single Ca^2+^ pulse with a speed given in Eq. 27 can traverse along an entire microtubule within the axoneme during a chlamydomonas typical beat period.

Otherwise, the relaxation time of a chlamydomonas pair of doublets considered as a slender beam of length *L* = 12*µm* with bending stiffness *κ* = 12*·*10^*-*24^ *Nm*^2^ and drag coefficient 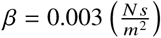 reads

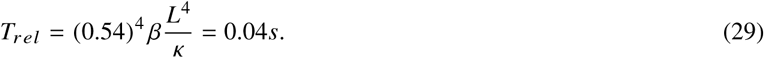

This indicates that an ionic pulse, Eq. 23 has enough time to activate motors prior to the bend being being relaxed.

### MODEL IN WHICH AXONEME CURVATURE REGULATES THE ACTIVITY OF DYNEIN MOTORS

Sartori et al. (11) presented how dynaimc axoneme curvature regulation plays a central role in tuning beats of chlamydomonas flagella. The flagellar beat is performed by dynein motors, which generate sliding forces between adjacent doublets.

Here, we rely on the experimental evidences concerning the partially disintegrated chlamydomonas axonemes published by Mukundan et al. (31) and Aoyama and Kamiya (32). They studied the interaction of pairs of MT doublets from partially split axoneme. Their assay has shown that these doublets were bent into circular arcs with an almost constant curvature.

The two doublets with constant spacing (*a*_0_) are constrained at the axoneme base (*ab*) with basal stiffness (2*K*_0_) of each doublet. Dynein act by producing a local sliding displacement of two doublets Δ(*s*) depending on the arc length position (*s*). The corresponding tangent angle is *ψ* (*s*). The normal *K*_*n*_ and shear stiffness *K* are broght about by the nexins of each axoneme. The MTs within doublets exhibit bending rigidity which was estimated in (31) to be

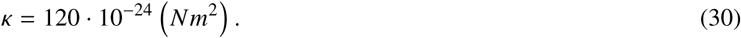

The dyneins step toward the base of doublets. This produces the force density on the top filament (1), + *f* (*s*) by subjecting it to tension and sliding it to the distal end. Conversely, the bottom filament (2) is under the compression with the same intensity-*f* (*s*). The mechanical energy of the filament-motor system is given by the following integral along the filament length *L*:

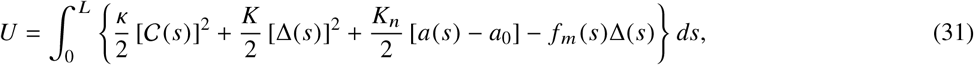

where the curvature of doublets *C*(*s*) is defined through the tangent angle *ψ* (*s*) with following derivative

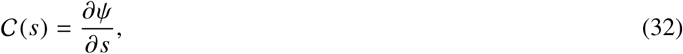

*a*(*s*) is the variable inter doublet spacing and *f*_*m*_ (*s*) is the total motor force per unit length of the doublet. The straightforward variation of the total energy *U* with respect to the tangent angle was elaborated in (31) and it yields

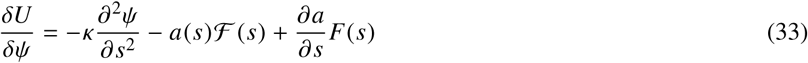

Here ℱ(*s*) represents the shear force per unit length

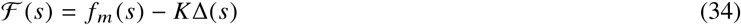

and the total shear force *F* (*s*) reads

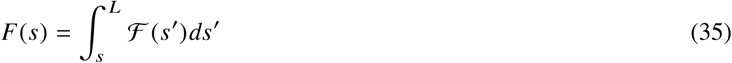

The variation of functional Eq. 31 with respect to separation *a*, for stationary condition gives:

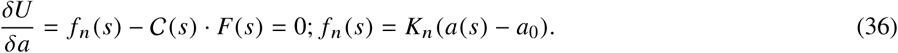

Based on the above, we now obtain the expression for normal force per doublet’s unit length as

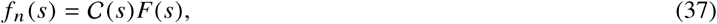

indicating that normal force depends on the total shear force and doublet curvature. This exhibit the mutual interplay between all three mechanisms involved in axoneme dynamics.

In this approach the inter-doublet spacing *a*(*s*) is considered to be a constant value *a*_0_ so that the normal force *f*_*n*_ (*s*) plays the role of a Lagrange multiplier.

Integrating Eq. 33 around the stationary state 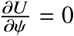 with a boundary condition of free doublet and *C*(L) = 0 and with the torque balance at base (*s* = 0)

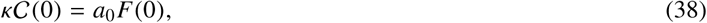

one obtains the value of the Lagrange multiplier as follows

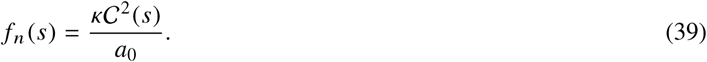

It expresses the nonlinear relation between normal force and doublet curvature. At a stationary configuration 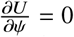, along with the fixed inter-doublet spacing *a*(*s*) = *a*_0_, from Eq. 33 we get the most important relation for the next section in the context of dynein regulation by the curvature control, namely:

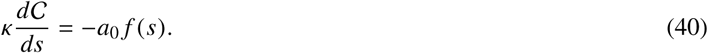

### THE ROLE OF Ca^2+^ IONS IN INITIATION AND REGULATION OF AXONEME BEATING

We now elaborate on the main point of this article. This is the mechanism, which involves the influence of Ca^2+^ ions in the initiation of axoneme beats through regulation of dynein motors activity.

We have shown that the polyelectrolite character of MTs provides a necessary condition for establishing the bell-shaped soliton-like pulses of cations, including the Ca^2+^ ions. These pulses can be injected through conformational changes of the Ca^2+^ conducting channels or exchangers.

Let one considers a 10-*µm* long cylindrical cilium with a diameter of 0.3*µm* yielding a 9.56*µm*^2^ surface area and a 0.94 *µm*^3^ = 0.94 *fl* volume, which is four orders of magnitude smaller than that of a typical cell. In comparison with such a cell itself the cilium has a 1.5-fold larger surface area-to-volume ratio. It implies that the same surface density of ion channels should change the ionic concentration within this small cilium volume remarkably faster than it occurs in the volume of a cell.

In that respect, in primary cilium (for example) the resting concentration of Ca^2+^ ions is seven-fold (0.7*µm*) greater than in cellular cytoplasm (0.1*µm*), Delling et al. (33). A very important issue is how beating cilia in (6a) the node drive left-right asymmetry in the vertebrate embrio (Babu and Roy, (34)). It appears that Ca^2+^ signaling in the node is much more dynamic than previously recognized. Takao et al. (35) puts forward the idea that protein PKD2 regulates the frequency of Ca^2+^ signals so that the frequency and not just the spacial distribution of Ca^2+^ ions is critical for initiating left-right asymmetry in an embrio. PKD2 is a 6-pass trans-membrane protein of 968 amino acids, which regulates the influx of Ca^2+^ ions into cilia axonemes. Alternatively, the flagellum may be sensitive to the rate of change of Ca^2+^ concentration increasing its beating asymmetry only in response to the rapid elevation in Ca^2+^ concentration produced by activation of the nimodipine sensitive pathway, Wood et al.(36).

Finally, in mammalian sperm, Ca^2+^ currents were only detected in the CatSper channels. CatSper is a sperm specific Ca^2+^ permeable, pH-sensitive and weakly voltage-dependent ion channel. It is located in the plasma membrane of the flagellar principal part (Sun et al. (37)). Besides the presence of other ionic channels only the CatSper directly modulates the physiological processes of sperm hyper-activation, sperm capaciatation, chemotaxis toward the egg and the acrosome reaction.

It is of particular importance to mention which parts of dynein are sensors for Ca^2+^ activation of motor binding to MTs.

Dynein of an outer arm in the axoneme contains three dynein heavy chains *α, β, γ* (DHCs). ATP-sensitive MT-binding by a dynein containing only *β* and *γ* DHCs does not occur at Ca^2+^ concentrations below pCa6 but is maximally activated above pCa5 (10*µM*).

These findings strongly suggest that Ca^2+^ binding directly to a component of the dynein complex regulates ATP-sensitive interactions between the *β* or *γ*-DHC and MT.

The observations by Sakato and King (38) indicate that the outer dynein arm contains a Ca^2+^ sensors that is responsible for flagellar waveform conversion during the photophobic response. They identified the light chain 4 (LC4) protein as the only Ca^2+^ binding component that is directly associated with outer arm DHCs. They also verified that LC4 acts as a sensor to mediate Ca^2+^ (but not Mg^2+^) dependent regulation of outer arm dynein motor function.

All the above suggests that *β* and/or *γ* DHCs with an associated LC4 protein play a pivotal role in Ca^2+^ regulation of flagellar motility.

It is important to note that the nexin link, the central pair-radial spoke and dyneins all contain proteins that bind Ca^2+^. Thus Ca^2+^ may indirectly regulate dynein activities by binding to multiple-bridging structures and thus modulating inter-MT spacing, Kamiya and Yagi (39). On the basis of afore-mentioned evidence we argue that the influx of Ca^2+^ ions is farther guided by MTs within an axoneme, thus being faster and more efficiently distributed towards the associated dynein motors leading to their activation in the process of binding to MTs.

Let us make the quantitative assessment of the role of solitonic ionic pulses sliding along MTs in axoneme dynamics.

Although electron microscopy studies have shown that there are longitudinal variations in dynein isoforms and dynein density, Bui et al. (40) we assume that the linear density of the same dynein isoforms *ρ* is uniform along doublets. A single attached motor exerts on doublet a positive force *f >* 0.

Motors attach to a doublet due to the activation initiated by Ca^2+^ ions and detach from the doublet due to its curvature and/or normal force. The probability that a motor at position *s* is being attached is *p*(*s*). Thus, the net motor force density can be expresses as follows

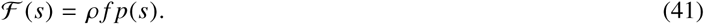

For a stationary configuration the probability *p* (*s*) reads, Bell (41):

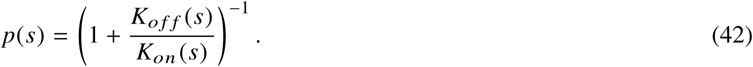

Smith (12) revealed that dynein activity increases in a linear manner with elevation of Ca^2+^ concentration. We here postulate that the attachment rate *K*_*on*_ (*s*) is being activated by Ca^2+^ ions, which are carried and distributed along MTs by a bell-shaped pulse described by the function following from Eq. 22

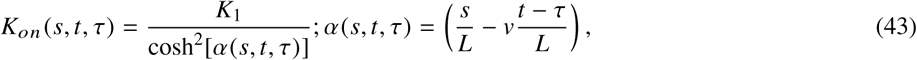

with *K*_1_ meaning the maximum of attachment rate and *τ* being the time delay caused by chemical inertia of the process. There is a delay between the action of the Ca^2+^ pulse and the response of dynein motors, Howard (6).

The maximum of the attachment rate is on the order of a characteristic ATP cycling rate

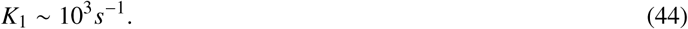

According to the experimental assay (31) the detachment of motors does comply to the curvature control mechanism in accordance with the fact that the detachment rate follows the Bell’s law (41):

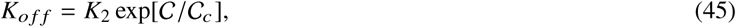

where *C*_*c*_ represents the characteristic crytical curvature indicating the onset of a significant increase of motor detachment.

In the case of Chlamydomonas flagella it amounts to

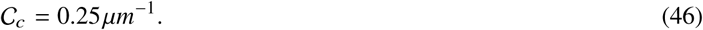

Now we can use Eq. 40 combining with Eqs. 41, 42, 43 and 44 to obtain

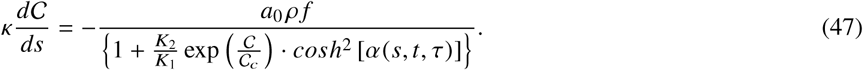

Following the experimental findings in (31), where for the most part in doublets the motors are detached indicating that condition (*K*_2_ *≥ K*_1_) holds, then in Eq. 47 the unity in the denominator can be safely disregarded yielding:

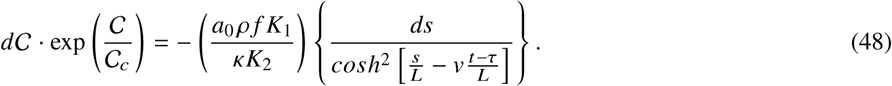

Using the abreviation

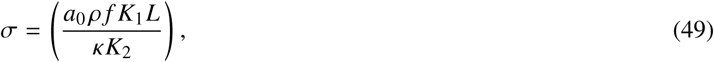

Eq. 48 could be readily integrated giving

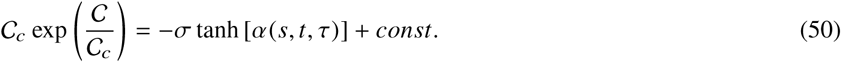

Using the boundary condition *C*(*L*) = 0, one obtains the explicit solution

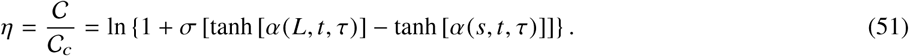

On the basis of the set of numerical parameters for chlamydomonas axoneme

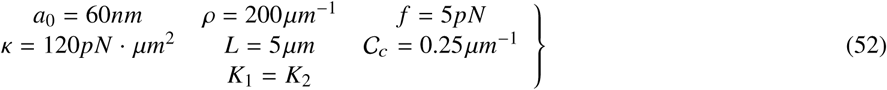

we find that the dimensionless parameter *s* amounts to

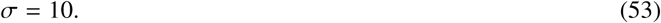

Eventually Eq. 51 for fixed (*t - τ*) has the dimensionless shape

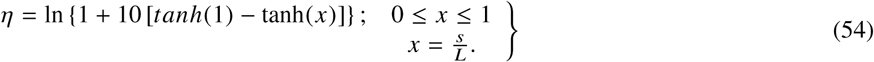

The maximal curvature appears for *x* = 0 with tanh 1 = 0.764 thus giving

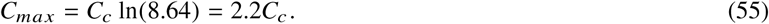

Sartori et al. (42) state that dynein motors in an axoneme respond to the time derivative of curvature rather than curvature itself. According to that concept we seek the derivative of Eq. 51 with respect to delay time *τ*

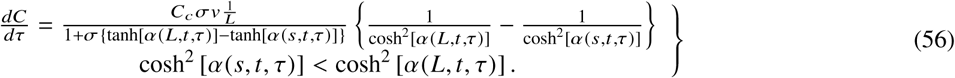

It appears that the condition

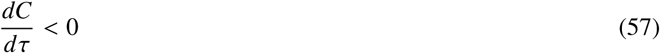

safely holds meaning that it agrees with the general theoretical approach developed in Sartori et al. (42).

It is meningful that the above derivative increases linerly with the speed of the solitonic ionic pulse. The maximum value of curvature time derivative, Eq. 56 follows for *s* = 0, *t - τ* = 0 thus giving

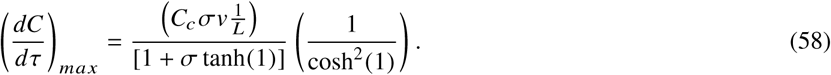

Inserting parameters for Chlamydomonas as 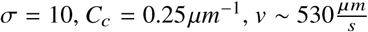 and *L*=5*µm*, one obtains

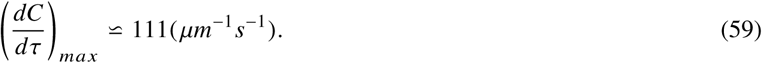

Finally, a few words about the role of post-translational modifications in CTTs of MTs within the axoneme. It appears that polyglutamination critically affects the function of inner arm dyneins. In cilia more than 10 glutamates are common reaching up to 28, Redeker (28).

It was demonstrated, Li et al. (43) that long-range electrostatic interactions between positive segments of dynein and negative CTTs of MTs bring a level of precision to an otherwise noisy dynein stepping process.

It seems that tubulin with highly acidic (negatively charged) polyglutaminated side chains elicits stronger attractive forces on dynein-e and dynein regulatory complex (DRC). Interestingly the dynein-e has the most positively charged MT-binding site among all DRCs, and is the most strongly attracted to MTs in this respect.

Regarding our model of Ca^2+^ pulses along MTs, on the basis of Eq. 5 it is clear that branched polyglutaminated CTTs have increased capacitance, so that a characteristic discharging time, Eq. 13 is being changed thus controlling the characteristic speed, Eq. 14. These changes in turn affect the curvature time derivative, Eq. 56, which is responsible for axoneme dynamics itself.

## DISCUSSION

The main contribution of this article is our original concept of how calcium ions control the beating dynamics of cilia and flagella by tuning the activities of dynein motors in axoneme. The crucial property in this process is the polyelectrolite character of microtubules within axoneme, which implicates the properties of nonlinear transmission lines. It enables positive ions including Ca^2+^ to form movable bell-shaped clouds around microtubules. The speed and stability of these ionic pulses enable a more precise and efficient mechanism for Ca^2+^ distribution compared with diffusion itself.

Experimental evidences, Smith (12) have shown that dynein activities in axoneme increased in a linear manner with an increasing Ca^2+^ concentration. However, an increasing Ca^2+^ concentration causes somewhat abruptly a switch from an asymmetric to a symmetric beat wave form. In the context of our model, at lower concentrations (smaller and slower ionic pulses) the binding of Ca^2+^ to LC4 and *β/γ* DHCs causes increasing activities of these dynein arms.

An additional increase of Ca^2+^ concentration (in terms of bigger and faster ionic pulses) brings about the neutralization of negatively charged polyglumatinated CTT chains. It causes the weakening of motor-MT interactions deactivating the dynein-arms within an associated bell-shaped Ca^2+^ pulse.

An important outcome of our approach is the fact that curvature control and curvature time derivative control mechanisms convincingly interplay with localized Ca^2+^ pulses enabling an almost quantitative agreement with experimental evidences for Chlamydomonas flagella, Mukundan (31).

Otherwise, the curvature control mechanism is unable to explain how dyneins could sense the very small tubulin strains associated with the observed curvatures (31). We suggest that instead of tubulin body the pertaining CTTs are the sensors for curvature control of dynein activities. This also arises from the fact that polyglutaminated CTTs sense dyneins in a different way compared with deglutaminated ones. This offers a possible solution to the problem of sensing the tiny strains associated with MT bending.

Recently Sartori et al. (42) showed that an axoneme twist provides an alternative mechanism by which the dynein can sense the curvature thus exhibiting the interplay between all three mechanism involved in axoneme dynaimcs (sliding, normal force and curvature control).

## ABBREVIATIONS

ATP: adenosine threephosphat
DHC: dynein havy chain
DRC: dynein regulatory complex
LC4: light chain 4
MT-microtubule: CTT-tubulin C-terminus tail

## AUTHOR CONTRIBUTIONS

MS and JT participated in basic idea and conception, analysis and interpretation of results. TN participated in design and drafting of article. DS participated in analysis and interpretation of data especially regarding concept of nonlinear transmission line.

## ACKNOWLEDGEMENTS

This research was financially supported by the Provincial Secretariat for Higher Education and Scientific Research of AP Vojvodina (Project No. 1144512708/201603), also by the Ministry of Education, Science and Technological Development of the Republic of Serbia (Project No. OI171009, III43008, and III45010), and by Serbian Academy of Sciences and Arts. J. A. T. ackowledges funding from MSERC (Canada).

## APPENDIX

In order to solve Eq. 20 we introduce a new auxiliary variable *y*(*ξ*)

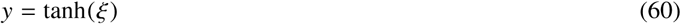

which gives the derivatives

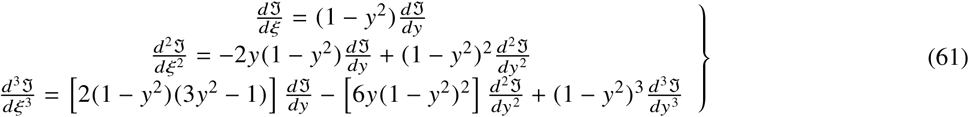

Inserting Eqs. 61 into Eq. 20 one obtains

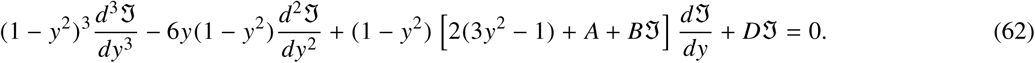

The unknown function 𝔍 (*y*) can be expanded in the power series:

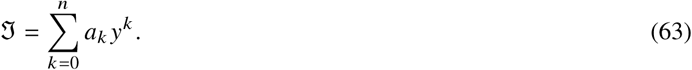

The idea is to make balancing of the order of nonlinear term with the dispersive term of third derivative in Eq. 62

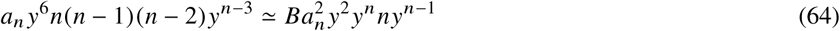

Or

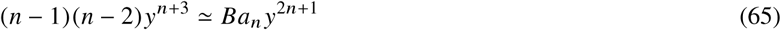

which yields

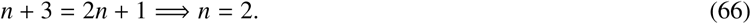

Thus the series, Eq. 63 reduces to the second power

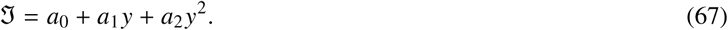

Taking the boundary conditions one obtains

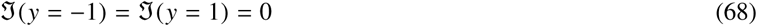

one obtains

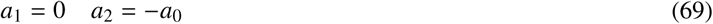

so that Eq. 67 takes the form

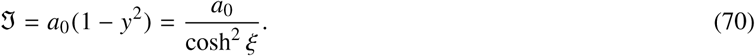

Inserting the function Eq. 70 and its derivatives in Eq. 62 we obtained

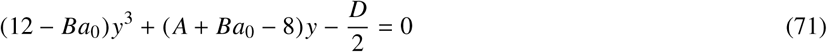

which implies

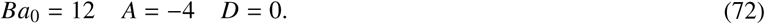

These relations lead to set of equations, Eq. 24.

